# Systematic mutational analysis of epitope-grafted ED3’s immunogenicity reveals a DENV3-DENV4 bi-serospecific ED3 mutant

**DOI:** 10.1101/2020.04.30.070250

**Authors:** Mamtaz Sultana, Nazmul Hasan, Mamunur R. Mahib, Manjiri R. Kulkarni, Yutaka Kuroda, Mohammad M. Islam

**Author notes:** Correspondence to: MMI,: YK.

## Abstract

Dengue viruses are classified into four serotypes (DENV1∼4), and the severe forms of dengue disease, the dengue hemorrhagic fever and shock syndrome, are caused by sero-cross-reacting antibodies. However, the residue determinants of the serospecificity and sero-cross-reactivity are yet to be identified. Here, we report an epitope grafting mutational analysis of the serospecificity and cross-serospecificity of the envelope protein domain 3 (ED3; 107 residues, ∼11.6kDa), which contains two major putative epitopes of DENVs. To this end, we constructed ED3 from DENV3 (3ED3) and DENV4 (4ED3), and six epitope-grafted variants, where we transferred epitope 1 (L^304^I, K^305^D, V^309^M, and S^310^A) and/or epitope 2 (D^383^N, K^384^S, K^387^T, and N^389^H) of 4ED3 onto 3ED3 and vice versa. Mice immunization using 3ED3 and 4ED3 generated serotype-specific antisera, as expected. Similarly, most epitope-grafted ED3s produced antisera serospecific to the template ED3 with little or no cross-recognition of ED3 of the serotype from which the epitopes were taken. This result indicated that a mere grafting of the epitope was not sufficient to transfer serospecificity, contrary to our expectations. However, one epitope grafted ED3 mutant, where epitope 1 of 3ED3 was grafted onto 4ED3 (4ED3^epi1^), generated antisera that was serospecific to both 4ED3 and 3ED3. The 4ED3^epi1^ is thus a chimeric ED3 that produces antisera possessing serospecificity to both 3ED3 and 4ED3. The 4ED3^epi1^ provides a unique tool for analyzing serospecificity and cross-reactivity in dengue, and we hope it will serve as a template for trivalent and eventually tetravalent antisera.

## Introduction

Dengue fever, a mosquito-borne viral disease, is caused by the dengue virus (DENV), which is a flavivirus classified into four serologically distinct serotypes (DENV1∼4). It is a major public health issue in tropical and subtropical regions [1,2], with 390 million cases reported every year, and 40% of the world’s population at risk [3,4]. Primary infection by a DENV may provoke a high fever for a few days, but the patient usually recovers and may gain a long-lasting immunity against the infecting serotype [5]. However, a secondary heterotypic infection can lead to severe syndromes such as dengue hemorrhagic fever (DHF) and dengue shock syndrome (DSS) [6,7]. The severity is thought to be caused by antibodies produced during the primary infection, which would be cross-reacting but sub-neutralizing against the secondary infecting DENV serotype. This phenomenon is coined as the Antibody Dependent Enhancement (ADE) [7–9]; ADE can be caused by natural dengue infection and also demonstrated in artificial immunization studies [7,10–12], which is a factor making the development of the dengue vaccine cumbersome [13,14]. The single-stranded RNA genome of DENV encodes ten gene products: The capsid (C), pre-membrane (prM), envelope (E), and seven nonstructural (NS) proteins [15]. The E-protein mediates virus-host attachment, and it is composed of three domains (ED1∼ED3) [16]. ED2, and accessorily ED3, are involved in the attachments of DENV to the host cell, and ED3 is the major target of neutralizing antibodies [17]. ED3s from all four DENV serotypes have a high sequence similarity (70-88%) and are structurally identical within an RMSD (Root mean square deviation) of 0.5-1Å [18]. Monoclonal antibodies against ED3 are neutralizing DENV in cell culture studies. Furthermore, the neutralization is corroborated by *in vivo* experiments in model mice [19-22]. Furthermore, immunization using ED3 can be long-lasting [23-25], suggesting that DENV’s ED3 is an essential factor in determining DENV’s serotype [26-29].

Despite numerous studies, the precise residue determinants of DENV serospecificity and sero-cross-reactivity are still to be fully identified [30-32]. In this study, we report the immunogenicity, serospecificity, and sero-cross-reactivity of ED3 from DENV3 (3ED3) and DENV4 (4ED3) and their epitope-grafted variants in Swiss albino mice. As expected, both 3ED3 and 4ED3 were serospecific: antisera recognized only the respective antigens. Further, the grafting of the putative epitope 1 and 2 individually or conjointly onto 3ED3 from 4ED3 and vice versa caused minor changes in the serospecificity of the antisera, which was against our initial expectation that serospecificity could be transferred by grafting the epitopes [19]. However, we observed that antisera raised against 4ED3 onto which epitope 1 from 3ED3 was grafted (4ED^epi1^), was serospecific to both 4ED3 and 3ED3. Thus, 4ED3^epi1^ generated a DENV3-DENV4 bivalent antisera, and we hope that it may provide a template for producing a tetravalent antiserum in the future.

## Results and Discussions

### Immunogenicity and serospecificity of 3ED3 and 4ED3

3ED3 and 4ED3 have a ∼70% sequence similarity, and their structures are very similar, if not identical (Figure 1) [18, 33]. However, despite being similar 3ED3 was slightly more immunogenic over 4ED3 in mice model (Figure 2; Suppl. Fig. S1-2) and produced highly specific antisera, in agreement with our previous report [26]. Namely, antisera raised against 3ED3 reacted only with 3ED3, and anti-4ED3 sera reacted only with 4ED3, with no or minimal cross-recognition (Figure 2; Suppl. Fig. S1-2). These observations demonstrated that even though ED3 is a fragment of the whole dengue envelope protein, it can generate a serospecific immune response with characters similar to those observed with the whole E-protein and the natural DENV [34,35].

**Figure 1.**
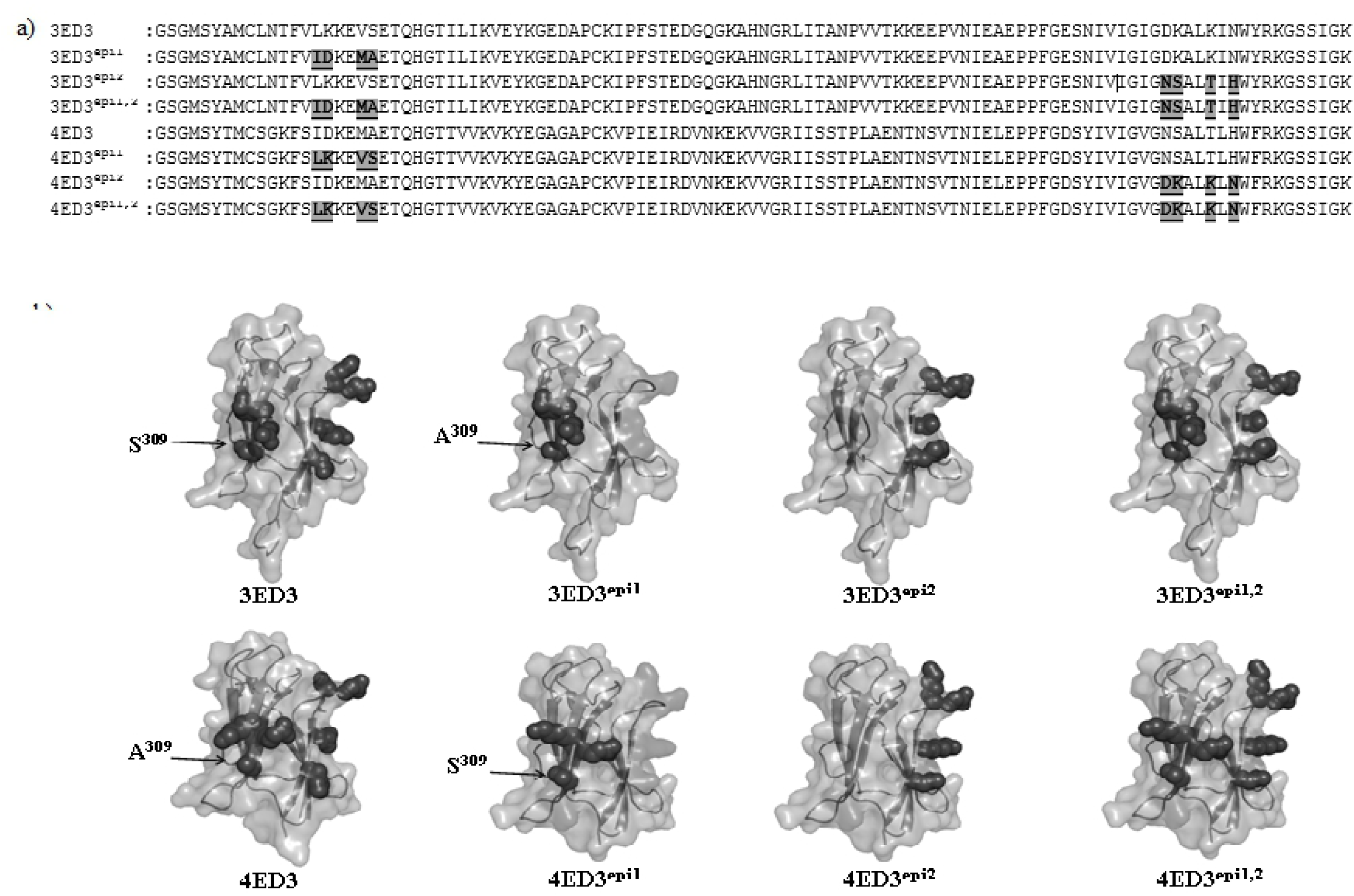
Sequences and structures of ED3s and their epitope-grafted variants. (a) Sequences of ED3 variants. Putative epitope residues grafted from 4ED3 onto 3ED3 and vice-versa are grey-shaded and underlined. (b) Surface model structures of 3ED3, epitope-grafted 3ED3 (3ED3^epi1^, 3ED3^epi2^, 3ED3^epi1,2^), 4ED3 and epitope-grafted 4ED3 (4ED3^epi1^, 4ED3^epi2^, 4ED3^epi1,2^) are shown in panel b. Structures of 3ED3 and 4ED3 were generated from 3vtt.pdb [18] and 3we1.pdb [33], respectively, while the surface structures of epitope-grafted ED3s were generated from their modeled structures (*per se* Materials and Methods). Grafted epitope residues on the protein surface are in black.

**Figure 2.**
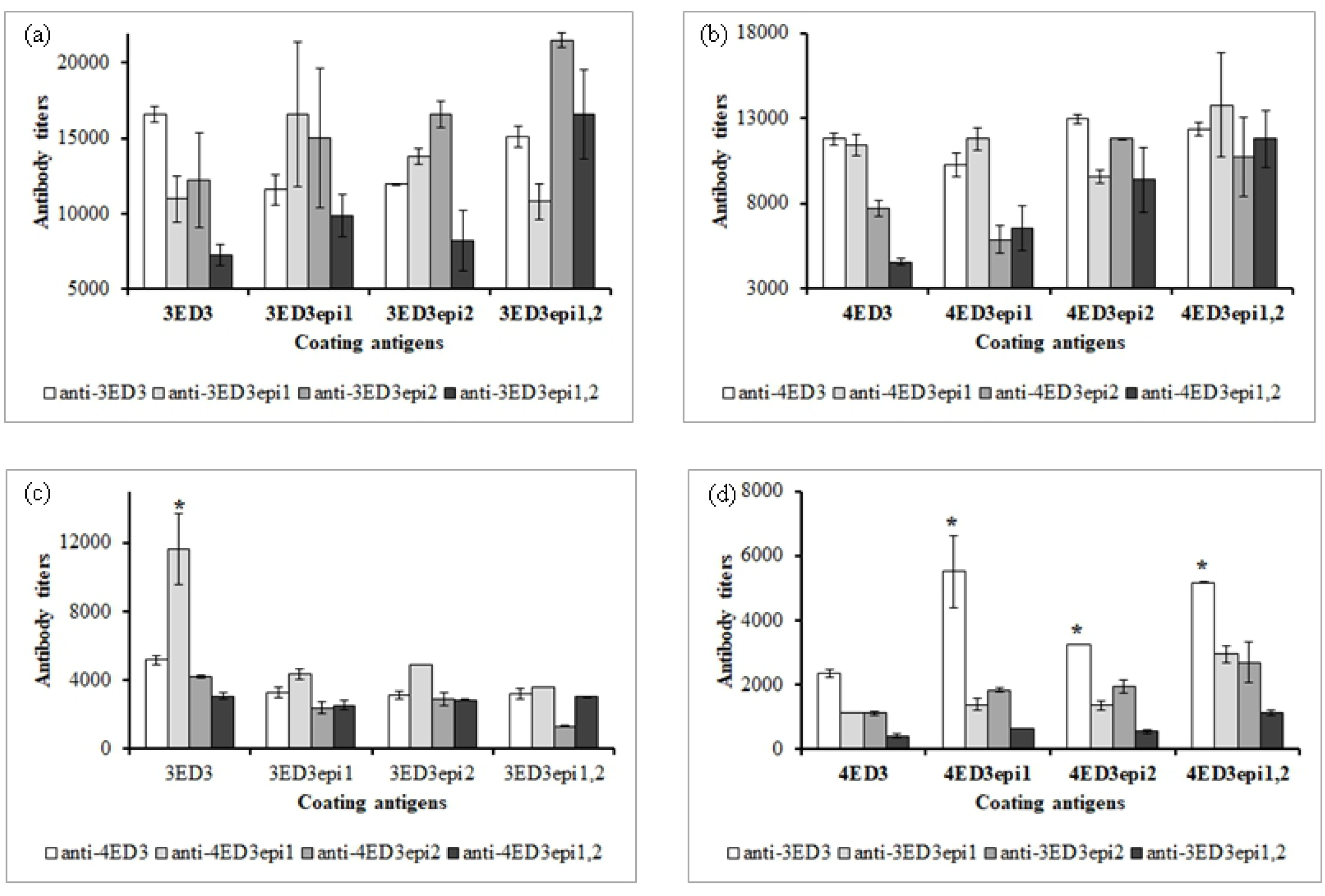
Effects of epitope-grafting on the serospecificity of 3ED3 and 4ED3. Antibody titers of anti-3ED3 and anti-epitope-grafted 3ED3 sera against 3ED3 variants (a), and anti-4ED3 and anti-epitope-grafted 4ED3 sera against 4ED3 variants. Cross-recognition of 3ED3 variants by anti-4ED3 sera (c) and cross-recognition of 4ED3 variants by anti-3ED3 sera (d) are shown.

### Effects of epitope-grafting on ED3s’ serospecificity

We grafted the putative epitope 1 (L^304^I, K^305^D, V^309^M, and S^310^A) and epitope 2 (D^383^N, K^384^S, K^387^T, and N^389^H) individually and conjointly from 4ED3 onto 3ED3, and vice-versa. We developed antisera against the two wild-type ED3s and their six epitope-grafted variants. The anti-3ED3 sera showed reduced recognition of epitope-grafted 3ED3 variants, where epitope 1 and 2 from 4ED3 were grafted onto the 3ED3 template (Figure 2a; Table 2; Suppl. Figure S1-3). Similarly, antisera raised against epitope-grafted ED3s exhibited the strongest recognition for the grafted 3ED3 variant against which was raised, and a reduced recognition for the wildtype 3ED3 (Figure 2a; Table 2; Suppl. Fig. S1a). In summary, these observations suggested that both epitopes play a major role in defining 3ED3’s serospecificity, since the recognition was lower when the epitopes were mutated.

**Table 1.** Putative epitope residues on 3ED3 and 4ED3.

| Epitopes | Residue position | Residue identities and ASA |  |  |  | Side-chain position |
| --- | --- | --- | --- | --- | --- | --- |
|  |  | 3ED3 (3vtt.pdb) |  | 4ED3 (3we1.pdb) |  |  |
|  |  | Identity | ASA (%) | Identity | ASA (%) |  |
| Epitope 1 | 304 | Leu | 23.5±4.2 | Ile | 17.6±1.8 | Buried |
|  | 305 | Lys | 40.5±2.6 | Asp | 36.8±1.8 | Partially buried |
|  | 308 | Val | 9.6±4.2 | Met | 5.9±2.6 | Buried |
|  | 309 | Ser | 67.4±1.3 | Ala | 80.2±8.0 | Surface |
| Epitope 2 | 383 | Asp | 83.4±13.0 | Asn | 85.3±0.5 | Surface |
|  | 384 | Lys | 76.0±5.7 | Ser | 42.3±9.6 | Surface/Partially Buried |
|  | 387 | Lys | 60.4±1.8 | Thr | 56.2±3.2 | Surface |
|  | 389 | Asn | 60.3±4.6 | His | 50.3±0.5 | Surface |

**Table 2.**
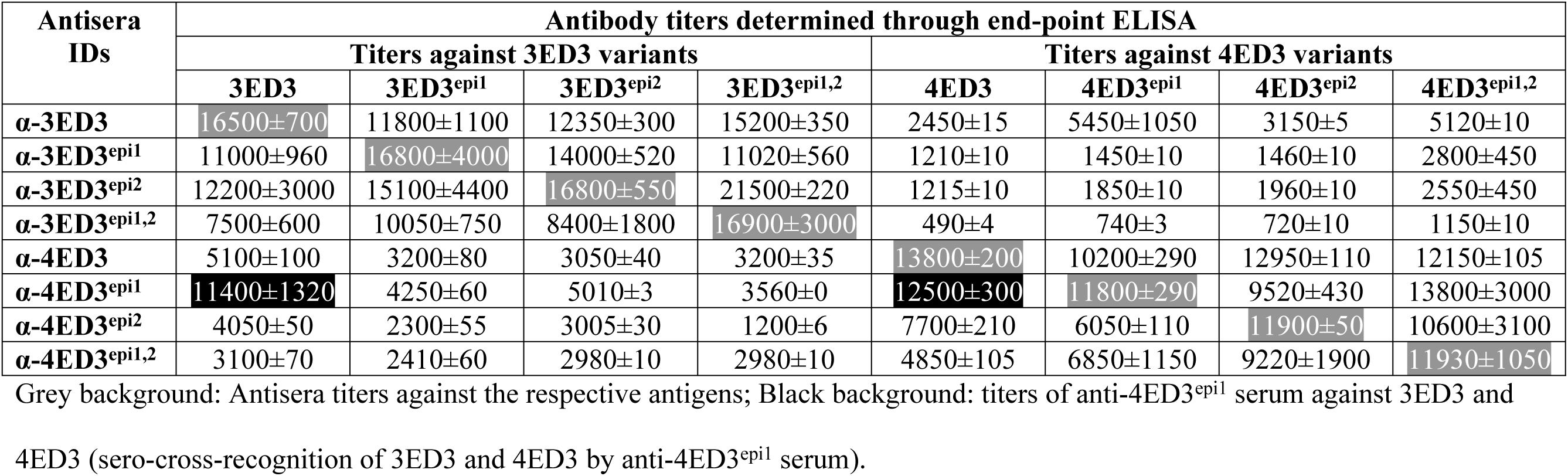
Antibody titers of anti-ED3 sera.

| Antisera<br>IDs | Antibody titers determined through end-point ELISA |  |  |  |  |  |  |  |
| --- | --- | --- | --- | --- | --- | --- | --- | --- |
|  | Titers against 3ED3 variants |  |  |  | Titers against 4ED3 variants |  |  |  |
|  | 3ED3 | 3ED3<sup>epi1</sup> | 3ED3<sup>epi2</sup> | 3ED3<sup>epi1,2</sup> | 4ED3 | 4ED3<sup>epi1</sup> | 4ED3<sup>epi2</sup> | 4ED3<sup>epi1,2</sup> |
| **α-3ED3** | 16500±700 | 11800±1100 | 12350±300 | 15200±350 | 2450±15 | 5450±1050 | 3150±5 | 5120±10 |
| **α-3ED3<sup>epi1</sup>** | 11000±960 | 16800±4000 | 14000±520 | 11020±560 | 1210±10 | 1450±10 | 1460±10 | 2800±450 |
| **α-3ED3<sup>epi2</sup>** | 12200±3000 | 15100±4400 | 16800±550 | 21500±220 | 1215±10 | 1850±10 | 1960±10 | 2550±450 |
| **α-3ED3<sup>epi1,2</sup>** | 7500±600 | 10050±750 | 8400±1800 | 16900±3000 | 490±4 | 740±3 | 720±10 | 1150±10 |
| **α-4ED3** | 5100±100 | 3200±80 | 3050±40 | 3200±35 | 13800±200 | 10200±290 | 12950±110 | 12150±105 |
| **α-4ED3<sup>epi1</sup>** | 11400±1320 | 4250±60 | 5010±3 | 3560±0 | 12500±300 | 11800±290 | 9520±430 | 13800±3000 |
| **α-4ED3<sup>epi2</sup>** | 4050±50 | 2300±55 | 3005±30 | 1200±6 | 7700±210 | 6050±110 | 11900±50 | 10600±3100 |
| **α-4ED3<sup>epi1,2</sup>** | 3100±70 | 2410±60 | 2980±10 | 2980±10 | 4850±105 | 6850±1150 | 9220±1900 | 11930±1050 |

Grey background: Antisera titers against the respective antigens; Black background: titers of anti-4ED3<sup>epi1</sup> serum against 3ED3 and
4ED3 (sero-cross-recognition of 3ED3 and 4ED3 by anti-4ED3<sup>epi1</sup> serum).

Corresponding experiments with anti-4ED3 and its epitope-grafted variants indicated a similar but not identical trend. Namely, anti-4ED3 sera titer against epitope-1-grafted 4ED3 (4ED3^epi1^) was slightly reduced while titers against epitope-2-grafted variants of 4ED3 remained almost unchanged (Figure 2b; Table 2; Suppl. Fig. S1b). However, anti-4ED3^epi1^ sera showed similar recognition of both 4ED3 and 4ED3^epi1^, suggesting that epitope 1 might not be a determinant for 4ED3 serospecificity. Interestingly, both anti-4ED3^epi2^ and anti-4ED3^epi1,2^ sera had very reduced recognition of wild-type 4ED3, indicating that epitope 2 might play a major role in determining 4ED3 serospecificity (Figure 2b; Table 2).

These observations indicated that the serospecificity transfer through epitope-grafting onto ED3s was scaffold-dependent. For example, we would conclude that epitopes 1 and 2 are essential for the serospecificity of 3ED3 because anti-3ED3 sera interaction with epitope-grafted 3ED3s was weak, and antisera raised against the epitope-grafted 3ED3 showed reduced recognition of 3ED3 (Figure 2a). However, antisera raised against 4ED3 sera exhibited a titer almost identical against all three epitope-grafted 4ED3s (Figure 2b).

Finally, let us consider sero-cross-recognition of 3ED3 and 4ED3 by antisera raised against the epitope-grafted ED3s. In most cases, no sero-cross-recognition was observed using epitope grafted mutants (Figure 2c-d; Table 2). The only exception was the anti-4ED3^epi1^ sera, which showed a significant recognition of 3ED3 (Figure 2c), in addition to recognizing 4ED3. This observation indicated that the serospecificity of DENV3 (3ED3) was almost entirely transferred onto DENV4 (4ED3) through grafting of epitope 1 (Figure 2d; Table 2). However, anti-epitope-grafted 3ED3 sera did show sero-cross-recognition of 3ED3 and 4ED3, again illustrating the fact that the serospecificity transfer was scaffold-dependent.

The identification of 4ED3^epi1^ as a chimeric ED3 possessing bi-serospecificity to both 3ED3 and 4ED3 is an important finding from an application viewpoint. This is because 4ED3^epi1^ may serve as a tool for further exploring serospecificity and sero-cross reaction among DENV serotypes. Furthermore, it might be used as a scaffold for producing a trivalent and, perhaps, a tetravalent antiserum against DENV.

### Structural origin of serospecific recognition of ED3s by anti-ED3s sera

For discussion, let us consider the structural origins of the serospecificity of ED3, based on the results of the grafting experiments. Grafting of the putative epitope 1 onto 3ED3 reduced serospecific recognition of anti-3ED3 sera (Figure 2). Among the four residues substituted, three were buried in 3ED3’s interior and were thus not expected to affect serospecific recognition of 3ED3 by anti-3ED3 sera [26]. The remaining S^309^A substitution was the only residue. Thus, it is tempting to consider that the S309A mutation was the main factor behind the reduced detection of anti-3ED3 sera by 3ED3^epi1^ and low antibody titers of anti-3ED3^epi1^ sera against 3ED3 (Figure 1; Table 1). However, this interpretation would need to be confirmed by single mutation analysis in the future.

Residues substituted on ED3s (residues grafted onto 3ED3 from 4ED3 and vice-versa); ASA: solvent Accessible Surface Area; ASA was calculated using 3vtt.pdb and 3we1.pdb; side-chain. ASA <30, 30-50, and >50% considered buried, partially buried, and surface-exposed, respectively.

On the other hand, all the four residues in epitope 2 are surface exposed and cause a significant modification in the size and properties of the side-chains (K^384^S, K^387^T, and N^389^H). Grafting of epitope 2 from 4ED3 onto 3ED3 template (3ED3^epi2^ and 3ED3^epi1,2^) induced noticeable local structural changes as assessed by circular dichroism (Figure 3; Suppl. Fig. S4). Such local significant structural changes may provide a rationale for the origin of the serospecificity.

**Figure 3.**
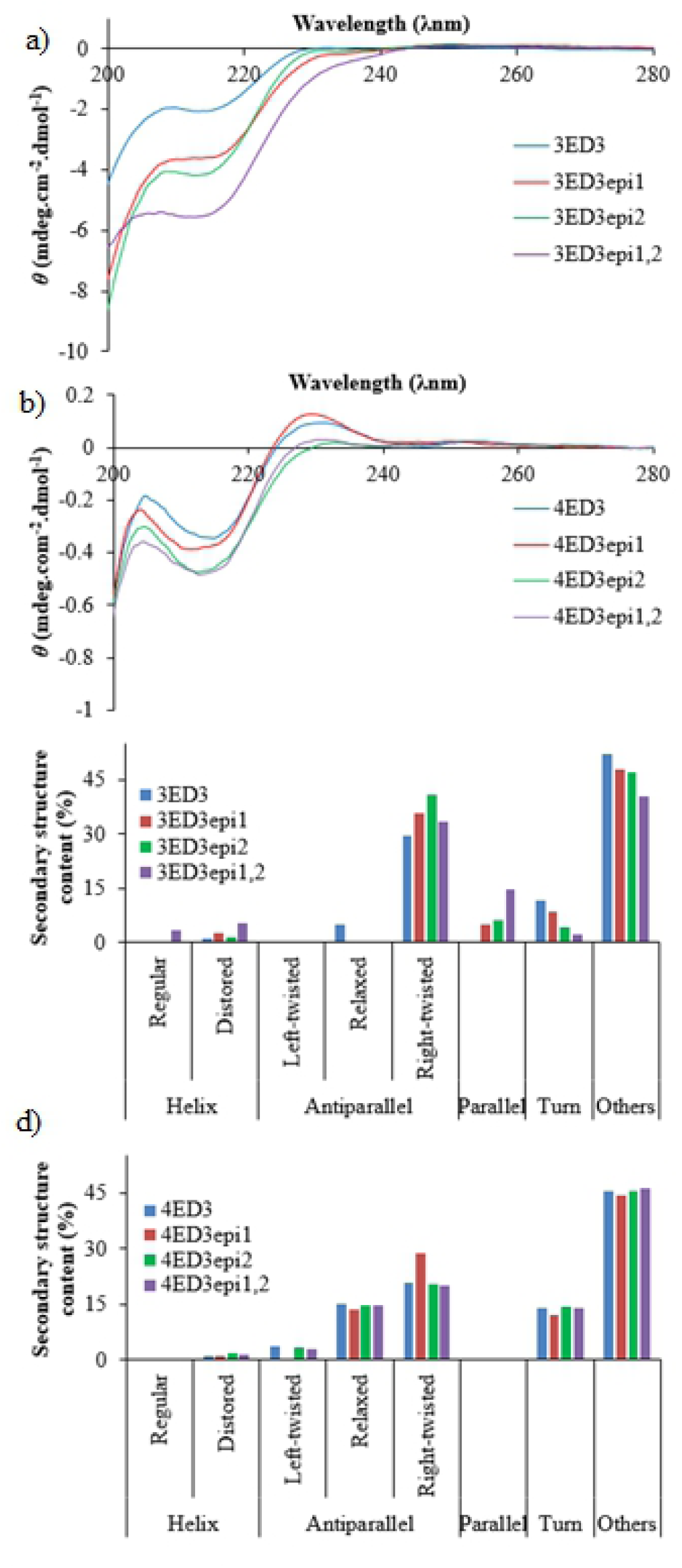
Effects of epitope-grafting on the structures of 3ED3 and 4ED3. (a) The CD-spectra of 3ED3 and its epitope-grafted variants and (b) 4ED3 and its epitope-grafted variants. CD-spectra were measured at 0.20mg/mL concentration in acetate buffer, pH4.5 at 30°C. CD spectra measured under immunization condition (at 0.2mg/mL in PBS are shown in Suppl. Fig. S4). The secondary structure contents of 3ED3 and 4ED3 variants estimated using bestsel [46] are shown in panel c, and d, respectively. Epitope-grafting on 3ED3 caused a noticeable change in the CD spectrum, but not on 4ED3. Legends are shown within the panels.

Furthermore, epitope-grafting from 3ED3 onto 4ED3 did not affect or barely affected the overall structure as assessed by the secondary structure content estimated from circular dichroism spectra data (Figure 3b-d) nor the serospecificity of 4ED3 (Figure 2). Besides, the CD spectra of 4ED3 and 4ED3^epi1^ remained almost unchanged under the immunization condition (at 0.2mg/mL in PBS; Suppl. Fig. S4b & d). This may explain why anti-4ED3^epi1^ sera had strong antibody titers against not only itself (and 4ED3^epi1^) but also against 3ED3, and 4ED3 (Figure 2; Table 2; Suppl. Fig. S4). The grafting of the putative epitope 2 barely or moderately affected the CD spectrum of 4ED3 [26], but it is yet unclear whether such small structural change is responsible for the loss of recognition (Table 2).

### Sero-cross-recognition of 3ED3 and 4ED3 by anti-4ED3^epi1^

As we discussed in the previous section, the substitution of epitope 1 on 3ED3 reduced recognition of anti-3ED3 sera and grafting of putative epitope 1 from 3ED3 onto 4ED3 significantly improved anti-3ED3 antisera’s recognition of 4ED3^epi1^ (Figure 2) without compromising the serospecific recognition of anti-4ED3 sera. The sero-cross-recognition of 4ED3^epi1^ by anti-3ED3 sera is attributed to a single substitution: Ser^309^ on 4ED3 (from 3ED3) (Suppl. Fig. S3). Furthermore, this interpretation is fully corroborated with the observation that antisera raised against 4ED3^epi1^ could equally recognize 3ED3, 4ED3, and 4ED3^epi1^, suggesting that Ser^309^ is responsible for the 4ED3^epi1^ chimeric ED3 detecting both anti-ED3s.

## Conclusion

In summary, this is most likely the first systematic examination of the effect of epitope grafting on the immune response and serospecificity of ED3 from DENV, which has four distinct serotypes. The results are not easily interpreted in terms of the serospecificity transfer, but we identified one mutant that holds a bivalent serospecificity against DENV3 and DENV4. The identification of 4ED3^epi1^ as a chimeric ED3 might act as a powerful tool for analyzing serospecificity and cross-reactivity in dengue; and we hope that it can provide a common scaffold that recognizes all four serotypes and possibly open the way for the design of a tetravalent dengue vaccine.

## Materials and Methods

### Mutant design, expression and purification

The sequences of DENV3 ED3 (3ED3) and DENV4 ED3 (4ED3) were retrieved from UniProt (ID P27915:1 and P09866, respectively) [36,37] and synthetic genes encoding ED3 sequences were cloned into a pET15b vector along with six His (His6-tag) and a thrombin cleavage site at the N-terminus, as reported previously [18,38]. The putative epitope residues (putative epitope 1: L^304^I, K^305^D, V^309^M, S^310^A; epitope2: D^383^N, K^384^S, K^387^T, and N^389^H) were grafted from 3ED3 onto 4ED3 and vice-versa through site-directed mutagenesis [26]. All ED3 variants were expressed in *Escherichia coli* JM109(DE3)pLysS as inclusion bodies and purified as previously reported [39]. Protein identities and purity were confirmed by analytical HPLC, and MALDI-TOF mass spectroscopy. The purified proteins were lyophilized and stored at −30°C until use.

### Circular dichroism (CD) measurement

Samples for CD spectra measurements were prepared at 5-10μM concentration in PBS, pH7.4 and acetate buffer, pH4.5, and centrifuged just before CD measurement to remove aggregates that might have accumulated during sample preparation [40]. CD spectra were measured using a 1mm cuvette with a JASCO J-820 spectropolarimeter in the wavelength range 200-260nm at different temperatures.

### Modeling of the epitope-grafted variants

The surface structure models for the epitope-grafted 3ED3 and 4ED3 were generated using the crystal structures of 3ED3 (3vtt.pdb) [18] and 4ED3 (3we1.pdb) [33], respectively, as templates. In short, we added the target epitope residues (see Figure 1a) to the template structures using COOT [41], assuming that the backbone structures remained unchanged. The side-chains were manually configured based on Richardson’s penultimate rotamer library [42] with no and/or minimal side-chain steric clashes as judged by Molprobity [43]. The surface structure models were generated using Pymol graphics [44].

### Artificial Immunization Studies

The artificial immunization studies were carried out in Swiss albino mice of 3-4 weeks of age (age at the start of immunization). Mice were repeatedly immunized with wild-type ED3s and their epitope-grafted variants at weekly intervals for five times at 20μg/dose in 100μl of PBS (Phosphate buffered saline, pH7.4) plus 100μl of Freund’s Adjuvants, as previously reported [26]. The first dose was given subcutaneously in a complete Freund’s adjuvant, and doses 2-5 were injected intraperitoneally in incomplete Freund’s adjuvant. The dose-specific antibody response was monitored through tail bleeding after each round of immunization using ELISA (*per se* next section). After the 5^th^ dose, mice were sacrificed by cervical dislocation. Sera were prepared from heart-blood collected after sacrificing and were preserved at −30°C until use.

### Evaluation of serospecificity and sero-cross-reactivity of anti-ED3 sera

Antisera developed in Swiss albino mice were first tested for serospecificity through enzyme-linked immunosorbent assay (ELISA) against the respective ED3 used as antigens and then tested for sero-cross-recognition using the other ED3 variants as coating antigens. Namely, 96-well microtiter plates (Nunc™) were coated with 100µl/well of ED3s (wild-type ED3s and their epitope-grafted variants) in PBS at 1.5µg/ml concentration for overnight at room temperature. Unbound ED3s were washed out with PBS, and plates were blocked with 1% BSA in PBS for 45 minutes at 37°C. After washing with PBS, mouse anti-ED3 sera were applied at an initial dilution of 1:500 fold, followed by a 3-fold serial dilution in 0.1% BSA-PBS-Tween20 and incubated at 37°C for 2 hours [26]. After an extensive wash with PBS-0.05% Tween20, 100μL/well of anti-mouse IgG-peroxidase conjugate was added at 1:3000 dilution in 0.1% BSA-PBS-Tween20 and incubated at 37°C, for 90 minutes. Finally, after thorough washing with PBS-Tween, coloring was performed with 100µl/well of OPD substrate (0.4mg/mL in 50mM citrate buffer, pH4.5 supplemented with 4mM H_2_O_2_) for 20 minutes followed by absorbance measurement at 450nm using a Multiskan Ascent (Thermofisher) microplate reader. Antibody titers were calculated from power fitting of absorbance versus reciprocal of serum dilution using a cutoff of 0.1 (OD_450nm_) above the background values [45].

## Acknowledgments

We thank Dr. Montasir Elahi for designing ED3 constructs and all members of the Kuroda Laboratory for discussion and technical assistance. We thank the International Centre for Diarrheal Disease Research, Bangladesh (ICDDR, B), for providing the Swiss albino mice.

## Funding

This research was supported by a GARE (MOE, Bangladesh) project grant and a Chittagong University Revenue budget grant to MMI, a GIR (Global Innovation Research, TUAT, Japan) invitation to MMI (FY 2018), and a JSPS grant-in-aid for scientific research (KAKENHI, 15H04359, and 18H02385) to YK.

## Contribution

MMI and YK designed the project and wrote the manuscript; MS and NH performed the experiment; MRM optimized the immunization dose and protocol; MRK helped to express and purifying ED3 variants. All authors read and approved the manuscript.

## Abbreviations

DENV: dengue virus
3ED3: DENV3 envelope protein domain 3
4ED3: DENV4 envelope protein domain 3
3/4ED3^epi^: epitope-grafted 3/4ED3
ELISA: enzyme linked immunosorbent assay.

## Conflict of interest

No conflict of interest.

## Supplemental Information

**Suppl. Fig S1. Effects of epitope-grafting on the serospecificity of 3ED3.** (a) ELISA (OD_450nm_) of anti-3ED3, anti-3ED3^epi1^ and anti-4ED3 against wild-type 3ED3 (a), 4ED3 (b), 3ED3^epi1^ (c) and 4ED3^epi1^ (d) are shown. Both anti-3ED3 and anti-3ED3^epi1^ recognized wild-type 3ED3, but anti-4ED3 did not cross-recognize 3ED3. Similarly, only anti-4ED3 recognized 4ED3, indicating that wild-type ED3s were serospecific, and epitope-grafting onto 3ED3 from 4ED3 did not much affect 3ED3 serospecificity and transfer 3ED3 serospecificity onto 4ED3. Grafting of putative epitope 1 and 2 onto 3ED3 reduced its ability to detect anti-3ED3 antisera. Similarly, antisera raised against epitope-grafted 3ED3 variants had reduced recognition of wild-type 3ED3. Similarly, epitope-grafting slightly reduced recognition of anti-3ED3 by 3ED3^epi1^ but did not improve recognition of anti-4ED3 and 3ED3^epi1^. However, sero-cross-recognition of 4ED3^epi1^ by anti-3ED3 improved following epitope-grafting onto 4ED3 from 3ED3. These observations indicated that epitope-grafting from 4ED3 onto 3ED3 could transfer serospecificity of 4ED3 onto 3ED3, and interestingly, serospecificity of 3ED3 could be transferred onto 4ED3 by epitope-grafting.

**Suppl. Fig S2. Effects of epitope-grafting on serospecificity of 4ED3.** (a) ELISA (OD_450nm_) of anti-4ED3, anti-4ED3^epi1^ and anti-3ED3 against wild-type 3ED3 (a), 4ED3 (b), 3ED3^epi1^ (c) and 4ED3^epi1^ (d) are shown. Epitope-grafting from 4ED3 onto 3ED3 did not improve recognition of 3ED3 and serospecific recognition of 4ED3 by anti-4ED3^epi1^ sera. Similarly, anti-4ED3 sera did not recognize 3ED3^epi1^ as well. On the other hand, anti-3ED3 and anti-4ED3^epi1^ sera showed similar recognition of 4ED3^epi1^, in addition to recognizing 4ED3. These observations indicated that 4ED3^epi1^ possesses serospecificity of 3ED3, in addition to maintaining serospecificity of 4ED3 (Figure 2b) and clearly suggest that 4ED3^epi1^ is a chimeric ED3 possessing serospecificity of both 3ED3 and 4ED3 onto a common 4ED3 scaffold.

**Suppl. Fig S3. Chimeric ED3 conferring serospecificity of both 3ED3 and 4ED3.** Serospecificity and sero-cross-recognition of wild-type 3ED3s and their epitope-grafted variants by anti-3ED3, anti-3ED3^epi1^, anti-4ED3 and anti-4ED3^epi1^ sera. 4ED3^epi1^ maintained serospecificity of both 3ED3 and 4ED3 and could recognize both anti-3ED3 and anti-4ED3 sera. Comparative view of antibody titers of anti-ED3 sera against 3ED3, 4ED3, and 4ED3^epi1^. Anti-4ED3^epi1^ had almost the same titers against 3ED3, 4ED3, and 4ED3^epi1^, indicating that 4ED3^epi1^ is a chimeric ED3 possessing serospecificity of both 3ED3 and 4ED3, both in generating cross-reactive anti-ED3 antibodies and detecting both anti-3ED3 and anti-4ED3 antibodies. Superimposed structures of 3ED3 and 4ED3 (c) and 3ED3 and 4ED3^epi1^ (d) (dark gray: 3ED3; light gray: 4ED3). Residues substituted (grafting of epitope 1) are indicated. The overall structure of modeled 4ED3^epi1^ is very similar to that of 3ED3.

**Figure S4. Effects of epitope-grafting on the structures of 3ED3 and 4ED3.** (a) CD-spectra of 3ED3 and its epitope-grafted variants and (b) CD-spectra of 4ED3 and its epitope-grafted variants. CD-spectra were measured under immunization condition (at 0.2mg/mL in PBS) and the secondary structure contents in 3ED3 and 4ED3 variants estimated using bestsel [46] are shown in panel c, and d, respectively. Epitope-grafting on 3ED3 resulted in a considerable change in the CD spectrum, but not on 4ED3. Legends are shown within the panels.

